# Loss of the habenula neuromodulator Kisspeptin1 disrupts learning in larval zebrafish

**DOI:** 10.1101/080994

**Authors:** Charlotte Lupton, Mohini Sengupta, Ruey-Kuang Cheng, Joanne Chia, Vatsala Thirumalai, Suresh Jesuthasan

## Abstract

Learning how to actively avoid a predictable aversive stimulus involves two steps: recognizing the cue that predicts upcoming punishment, and learning a behavioral response that will lead to avoidance. In zebrafish, ventral habenula (vHb) neurons have been proposed to participate in both steps by encoding the expected aversiveness of a stimulus. vHb neurons increase their firing rate as expectation of punishment grows, but reduce their activity as avoidance learning occurs. How the change in vHb activity occurs is not known. Here, we ask whether the neuromodulator kisspeptin1, which is expressed in the ventral habenula, could be involved. *Kiss1* mutants were generated with Cas9 using guide RNAs targeted to the signal sequence. Mutants, which have a stop codon upstream of the kisspeptin1 peptide, have a deficiency in learning to avoid a shock that is predicted by light. Electrophysiology indicates that kisspeptin1 has a concentration-dependent effect on vHb neurons: depolarizing at low concentrations and hyperpolarizing at high concentrations. These data suggest that as the fish learns to cope with a threat, kisspeptin1 may differentially modulate vHb neurons. This implies that learning a behavioral strategy to overcome a stressor is accompanied by physiological change in habenula neurons.

**Significance statement:** Learning to deal with adversity can positively affect one’s ability to cope with challenges in the immediate future. Control thus causes short-term change in the brain. Here, we show that the neuromodulator kisspeptin1 is required to learn to avoid a punishment. Expression and electrophysiological recordings suggest that this molecule functions by controlling the ventral habenula, a region of the brain that mediates fear by regulating serotonin release. Kisspeptin1 may be a potential player in resilience developed as a result of control, extending previous findings that it can reduce fear.

## Introduction

When faced with an aversive stimulus, animals respond in a manner that is dependent on the context and experience. Upon first encountering such a threat in a novel environment, there may be panic and poorly directed attempts at escape. If they repeatedly encounter the threat, and become familiar with a safe escape route, they will be able to quickly remove themselves from danger. Better still, if the animals are able to recognize a cue that reliably predicts the impending threat, they will be able to escape before the aversive stimulus is present. This, in essence, is the phenomenon of active avoidance. Several theories have been proposed to explain the mechanism underlying active avoidance. In the two-factor theory, the animal first develops a fear of the conditioning stimulus (CS) that is paired with the aversive stimulus, by Pavlovian conditioning. Termination of the CS and the threat (unconditioned stimulus; US) then drives learning. Expectation has a critical role, and actions that lead to better than predicted outcomes are reinforced (1). The first time an aversive stimulus is encountered, the predicted outcome would be negative. However, if an escape route has been learnt, then the predicted outcome becomes positive.

How is active avoidance implemented in the brain? One structure that appears to be involved is the lateral habenula, which receives reward information from the globus pallidus of the basal ganglia (2). As first shown in monkeys, unexpected punishment leads to increased activity in lateral habenula neurons, while an unexpected reward leads to inhibition. In zebrafish, unexpected punishment leads to phasic activity in the ventral habenula (vHb) (3), which is the homolog of the mammalian lateral habenula. As the animal learns to associate a CS with the threat, there is tonic firing to the CS in vHb neurons. This causes excitation of serotonergic neurons in the dorsal raphe. As the animal learns to escape, there is decreased tonic firing. Tonic activity in the vHb has thus been proposed to encode aversive reward expectation value.

What is the mechanism of change in vHb activity as learning occurs? In zebrafish, Amo et al (3) have proposed that excitation is regulated via feedback from serotonergic neurons in the raphe to the entopeduncular nucleus, which is the teleost homolog of the basal ganglia. Whether additional mechanisms are involved is unknown. Here, we examine the possibility that a change in habenula neurons accompanies the learning process. In particular, we examine the potential involvement of the neuromodulator kisspeptin1, which is exclusively expressed in the vHb, together with its receptor (4). In zebrafish, two paralogs of *kiss1* have been identified. These have non-overlapping expression, with *kiss1* being restricted to the habenula, while *kiss2* is expressed in the hypothalamus and posterior tuberculum (4). This allows a specific test of the role of *kiss1* in the habenula via genetics.

In mammals, *kiss1* is expressed in the hypothalamus, and is well studied in the context of reproduction. Burst firing of hypothalamic neurons leads to the release of Kisspeptin1, which causes immediate depolarization of gonadotropin releasing hormone (GnRH) neurons. Kisspeptin1 is also expressed in the hippocampus, where it causes an increase in EPSC and contributes to increased excitability (5). In the zebrafish habenula, kisspeptin1 has been proposed to depolarize vHb neurons, based on c-fos expression (6). However, delivery of Kisspeptin1 decreases fear, which is inconsistent with evidence that excitation of vHb is aversive (3). Moreover, lesioning of Kisspeptin1 receptor-expressing neurons in ventral habenula neurons mimics the effect of Kisspeptin1 delivery (6), which would not be expected if Kisspeptin1 causes depolarization. Hence, although Kisspeptin1 is well placed to alter responses of the habenula to aversive stimuli, how it functions is unclear. Additionally, whether it is involved in instrumental learning is unknown. Here, we address both these questions.

## Results

As in adult zebrafish (7, 8), kisspeptin1 is present in the ventral habenula of larval zebrafish, together with its receptor, kiss1rb (Fig. 1A, B). The kisspeptin system is thus expressed in an appropriate manner to locally regulate vHb neurons even at an early stage. To test whether kisspeptin1 is required for avoidance learning, we generated mutations in the *kiss1* locus using CRISPR/Cas9. Two guide RNAs were designed to the signal peptide region of kisspeptin1 (Fig. 1C). These were injected into embryos at the 1-cell stage, together with mRNA for Cas9. High-resolution melt analysis of genomic DNA derived from injected embryos indicated that the guide RNAs were effective. This was confirmed by sequencing: 8/8 injected embryos contained mutations at the target site. Injected siblings were thus grown up. Sequencing of F1 fish indicated a transmission rate of 100%. Three alleles were obtained: one consisted of a 20 base pair deletion (kiss1^sq1sj^; Fig. 1D), while the other two were insertions of 7 and 8 base pairs (kiss1^sq2sj^ and kiss1^sq3sj^; Fig. 1D); kiss1^sq3sj^ contained a one base pair deletion, resulting in the same reading frame as kiss1sj^sq2sj^. All alleles gave rise to premature stop codons, upstream to the kisspeptin1 peptide (Fig. 1E). No label could not be detected in mutant fish by immunofluorescence with an antibody to the C-terminus of prepro-kisspeptin1 (7) (Fig. 1F), further indicating that the mutants lacked the peptide.

**Fig. 1.**
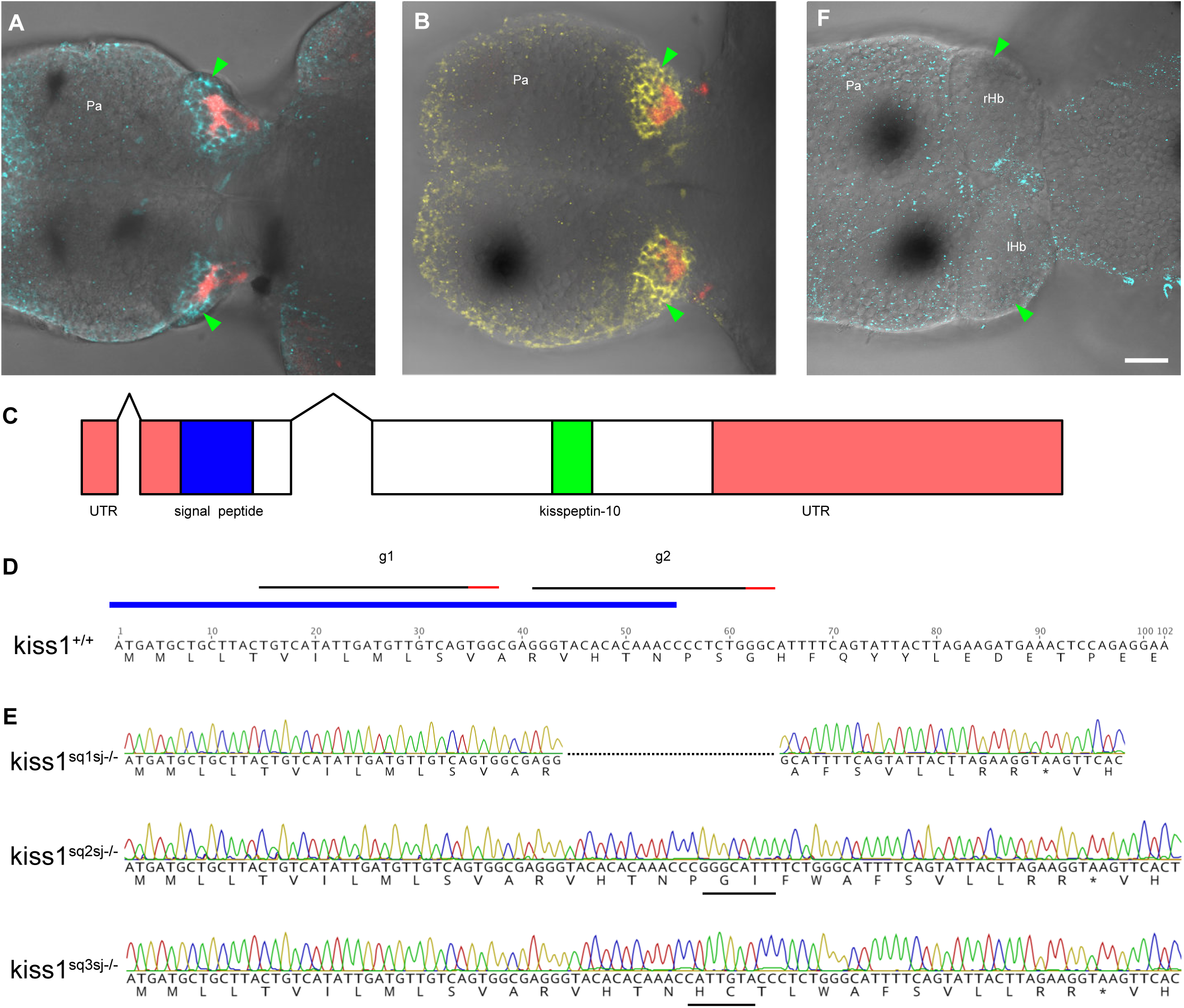
Generation of mutations affecting the zebrafish *kiss1* gene using CRISPR/Cas9. (A, B) Dorsal view of zebrafish larvae, labeled with an antibody to kisspeptin1 (A; cyan) and the kisspeptin receptor (B; yellow). These fish are from the *Et(SqKR11)*transgenic line, and express red fluorescence in afferents from the entopeduncular nucleus (basal ganglia) that terminate in the neuropil of the ventral habenula. The green arrowhead indicates the habenula. (C) A schematic of the *kiss1* gene. There are two introns. The signal sequence is shown in blue, and the region containing the mature kisspeptin1 peptide is shown in green. (D) Partial sequence of the *kiss1* gene. The signal sequence is indicated by the blue bar. The position of the two guide RNAs are indicated by the black bars. (E) The sequence of the three mutant alleles, together with predicted translations. The asterisks indicate stop codons. The black bars indicate inserted sequences. (F) A *kiss1^sq1sj -/-^* fish, following labeling with the kisspeptin1 antibody. No signal could be detected in the habenula. Scale bar = 25 µm. Pa: pallium; rHb: right habenula; lHb; left habenula.

To test their ability to learn to avoid an aversive stimulus, juvenile mutants and wildtype animals (5-6 weeks of age) were tested individually in a tank with two compartments (Fig. 2A). This is similar to a previously described apparatus (9), with the addition of a partial separator between the compartments as well as full automation (10). Each compartment contained a red light (the CS) and electrodes that delivered an aversive shock. The light was turned on for 8 seconds in the compartment containing the fish, and co-terminated with the shock if the fish still stayed in the CS side. If the fish moved to the non-CS side and stayed there until the end of the CS presentation, no shock was delivered. Fish were exposed to 10 training trials. The cross score was calculated from the number of times an individual fish swam to the non-CS chamber before the light was turned off. *Kiss1* mutants showed a lower cross score when compared to wild types (Fig. 2B), suggesting that kisspeptin1 is involved in learning active avoidance.

**Fig. 2.**
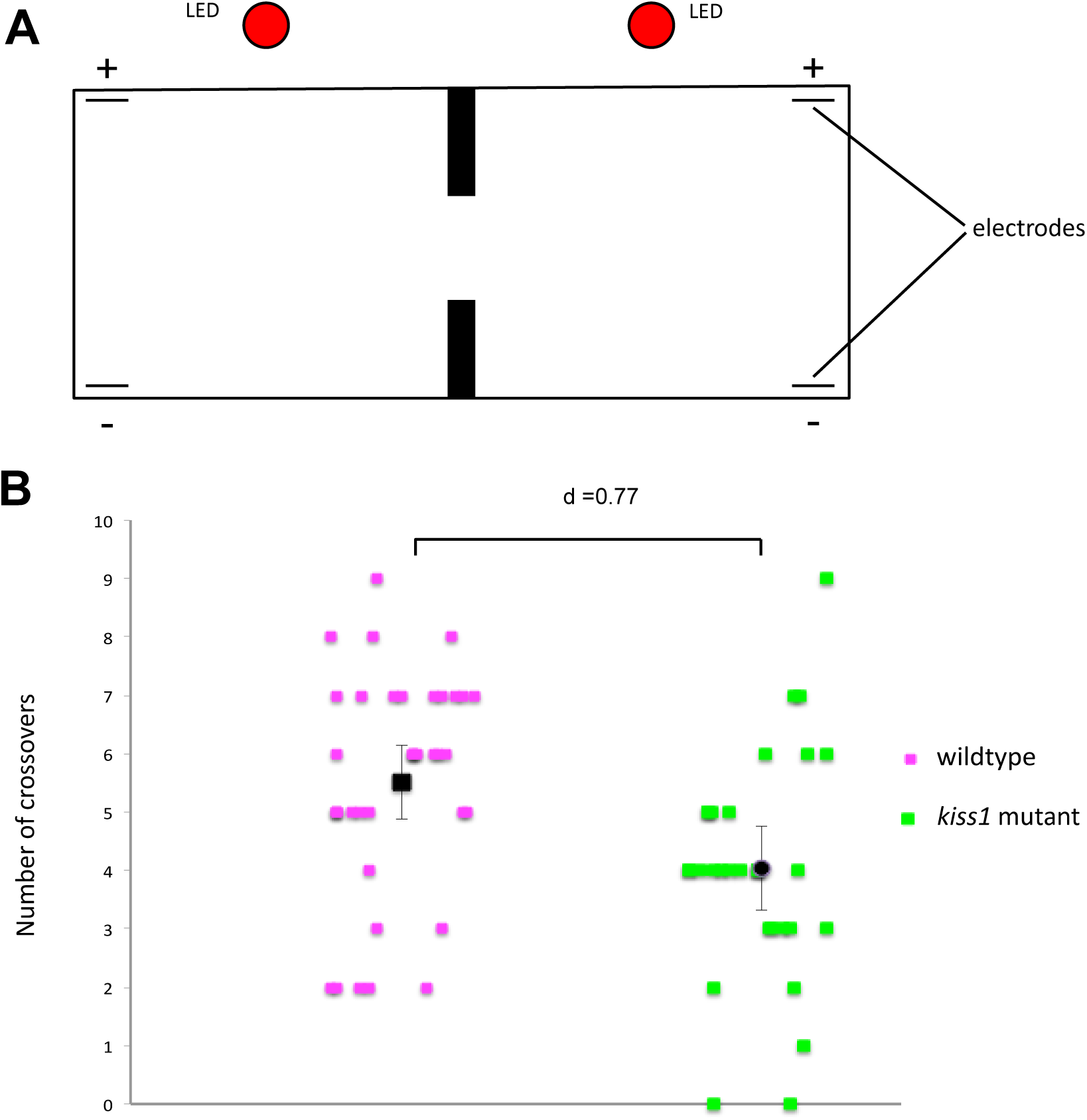
The behavior of *kiss1* mutants in an active avoidance assay. (A) Schematic of the two-way active avoidance chamber used. The fish were exposed to ten trials of light that was co-terminated with a shock. (B) The number of crossovers prior to shock delivery. The black symbols are average values, while error bars indicate 95% confidence interval. The performance of wildtype fish is different from mutants. Effect size, d = Cohen’s d. p = 0.0017 (Mann-Whitney U test, U = 319, Z = 3.13).

To determine how Kisspeptin1 affects habenula neurons, we performed whole cell patch clamp recordings from these cells and applied K-10, a conserved 10 amino acid peptide of Kisspeptin1. 1 µM TTX was added to the bath solution to globally block network activity. Cells were recorded in voltage clamp mode and were taken through a series of 500 ms voltage step protocol (Fig. 3A, lower panel). The cellular response (Fig. 3A, upper panel) was recorded before and after bath application of K-10. The recording was allowed to stabilize for 5 minutes in normal saline before K-10 was applied. Input resistances and holding currents did not change significantly with application of K-10 (p>0.05, SignTest). Difference currents were then calculated by subtracting the current values after K-10 application from the one before K-10 application (control). Difference currents calculated this way, showed that 5 µM K-10 application induced an outward current at depolarized potentials (Fig. 3B, n=4 cells from four 7dpf larvae). The mean peak amplitude of this current at 25 mV from four cells was 13.3 ± 8.9 pA. Next, to determine the specificity of this current, we bath applied a kisspeptin antagonist, 5 µM K-234 (11) along with 5 µM K-10, in a different set of experiments. The outward current was blocked in cells exposed to K-234 (Fig. 3D), confirming that this current was indeed induced specifically by K-10. To check whether kisspeptin1-evoked current exhibited a dose response, we repeated this experiment at 10 nM, 100 nM and 1 µM concentrations of K10. Lower concentrations of K-10 application appeared to induce lower amplitudes of the peak current at 25 mV (Fig. 3C), with 100 nM inducing an inward current, suggesting that low and high concentrations of kisspeptin1 have opposite effects on habenula neurons.

**Fig. 3.**
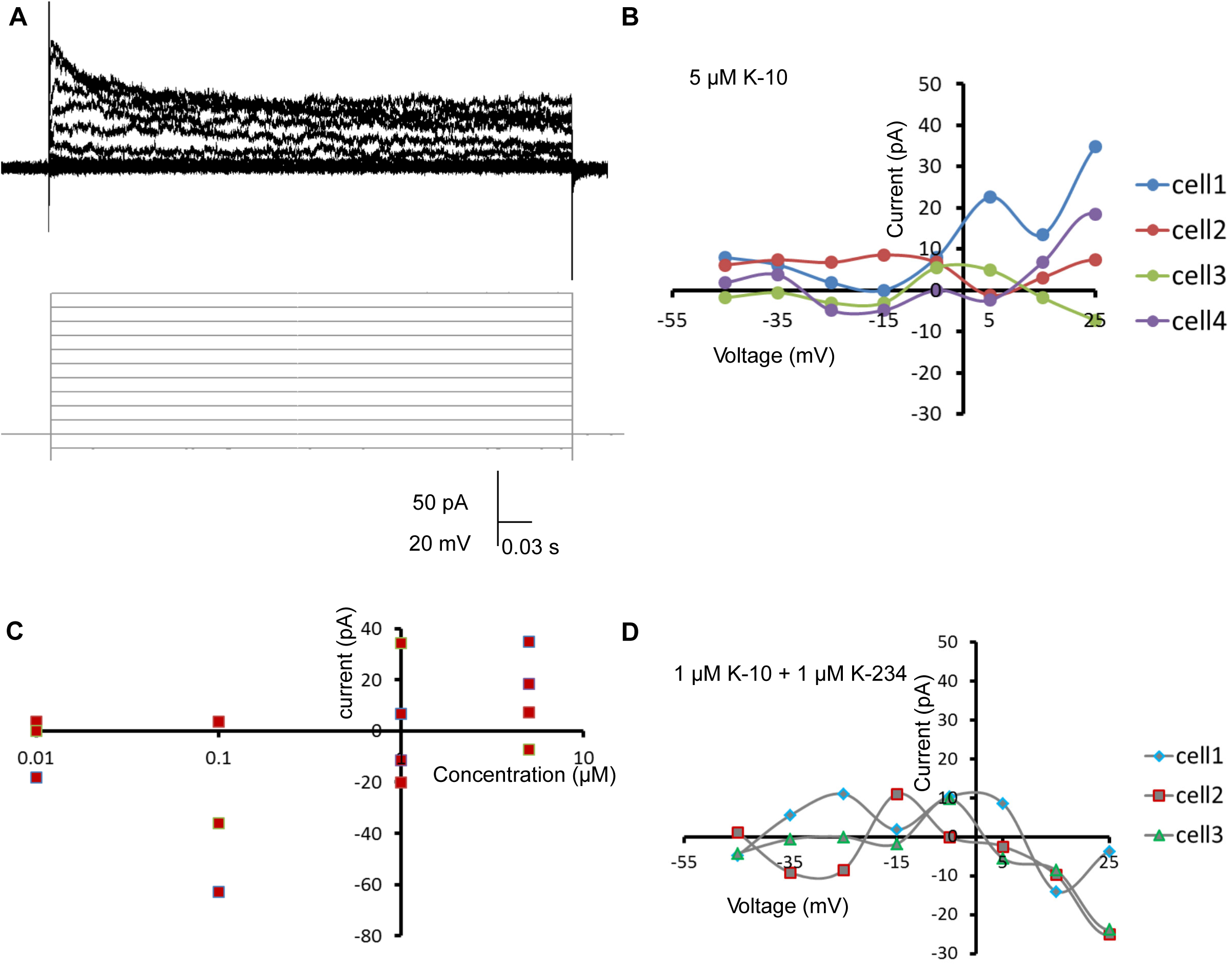
The effect of kisspeptin1 on vHb neurons. (A) Representative trace of cellular response (top, black) in a cell to voltage steps (bottom, grey). These recordings were done in the presence of 1 M TTX to block network activity. (B) Difference current obtained after bath application of 5 M K-10. The same protocol as in A was done before and after bath application of K10. The traces obtained after were subtracted from the traces obtained before application of the peptide for the same cell. (C) Dosage response of the difference current at 25 mV showing higher current response for higher concentrations of K-10, plotted for four different K-10 concentrations (10nM, 100nM, 1M and 5M). (D) Difference current plotted in the presence of the antagonist (1M K-234) and 1M K-10 showing the current response was inhibited in the presence of the antagonist.

## Discussion

We have investigated the role of the neuromodulator kisspeptin1 in active avoidance learning in zebrafish. Kisspeptin1 is expressed only in the habenula of zebrafish, primarily in ventral habenula neurons that project to the raphe (7). The kisspeptin1 receptor Kiss1rb is expressed in habenula neurons and not in downstream neurons. This, together with the defect seen in *kiss1* mutants, suggests that active avoidance learning involves kisspeptin1 signaling within the habenula. However, given that loss of Kiss1 did not completely eliminate learning in all fish, other mechanisms must also be involved. These may include a change in input from the basal ganglia (entopeduncular nucleus), via feedback from the raphe (3).

A function for kisspeptin1 in regulating fear responses was suggested previously based on the finding that injection of the peptide into brain ventricle of adult zebrafish led to depolarization of habenula neurons, as assessed by *c-fos* expression, and a reduction of innate fear (6). Surprisingly, destruction of cells containing the kisspeptin1 receptor, including vHb neurons, had the same effect as administering kisspeptin1 peptides. This raises a conundrum: how can stimulating a neuron have the same effect as killing the neuron? The present results provide one way to resolve this contradiction, which is that Kisspeptin1 has a concentration-dependent effect on ventral habenula neurons: low-concentrations lead to depolarization, while high concentrations lead to hyperpolarization. A concentration-dependent effect for kisspeptin has been reported before, for example in GnRH neurons of the medaka (12), with only low concentrations causing depolarization. Also, Ogawa et al reported that 10^−11^ mol/g body weight of kisspeptin1 increased *c-fos* expression in the habenula, whereas a higher concentration of 10^−9^ mol/g did not (13). Successful learning may be accompanied first by kisspeptin1-mediated excitation, and then by inhibition of vHb neurons. This would increase aversive expectation whilst the CS is being associated with US, and reduce aversive expectation as the strategy to avoid the US is learnt. What controls the release of kisspeptin1 as a function of learning is unclear.

Kisspeptin may modulate neurons in the mammalian habenula, given that the kisspeptin receptor is expressed in the lateral habenula of mice and rats (14, 15). However, the role of this expression has not been addressed. Intriguingly, experiments in mammals demonstrate that the habenula and the downstream raphe (16) have a role in the sustained effects of a stressor on subsequent response to challenges (17). It is tempting to speculate that kisspeptin may be involved in this phenomenon. For example, if the animal has learned active avoidance, kisspeptin1 may be at a level that reduces subsequent activation of the raphe. The finding that delivery of kisspeptin1 peptides blocks fear in zebrafish (6), in a serotonin-dependent manner (18), is consistent with this idea. It would be interesting to test the effect of delivery kisspeptin to the habenula of mammals, to see if high levels of this peptide can reduce the effects of uncontrollable stress.

## Materials and Methods

### Animals

Experiments were carried out on the AB strain of zebrafish, Danio rerio, in accordance with protocols approved by the Institutional Animal Care and Use Committee.

### Antibody labeling

Standard methods were used. Briefly, fish were fixed overnight at 4˚C, in 4% para-formaldehyde in PBS. A solution of PBS with 1% bovine serum albumin (Fraction V; Sigma), 1% DMSO and 0.1% Triton X-100 was the used to permeabalize the tissue and to dilute primary antibodies. The antibody to kisspeptin1 and the receptor kissr1b have been described previously (7, 8). Alexa488-conjugated goat anti-rabbit antibodies (Invitrogen) were used at 1:1000 dilution, in PBS. Imaging was carried out using a Zeiss LSM510 confocal microscope, with a 40x water immersion objective.

### Mutagenesis of the *kiss1* locus

Guide RNAs to the kiss1 gene were designed using ZiFiT Targeter (http://zifit.partners.org/ZiFiT/) (19). The Basic Local Alignment Search Tool (BLAST) was used to test for off-targets and only those target sites that yielded no identical off-targets were used. The chosen target sites were integrated into a forward primer (GAAATTAATACGACTCACTATAGGN_18_GTTTTAGAGCTAGAAATAGC (20)). PCR was performed with Phusion High-Fidelity polymerase (ThermoScientific) and a universal reverse primer that defined the remainder of the sgRNA sequence (AAAAGCACCGACTCGGTGCCACTTTTTCAAGTTGATAACGGACTAGCCTTATTTTAACTTGCTATTTCTAGCTCTAAC). PCR products were purified (QIAquick PCR purification kit, Qiagen) and 0.1ug was transcribed using the MEGAshortscript^TM^ T7 Transcription Kit (Life Technologies). sgRNAs were purified using ammonium acetate precipitation (2.5 µl 0.1M EDTA (Promega), 5 µl 5M ammonium acetate solution (Qiagen), 115 µl 100% ethanol) and stored in 1 µl aliquots at -80°C.

The Cas9 expression vectors pT3Ts-nls-zCas9-nls (Addgene plasmid #46757) (21) was linearized using XBaI (New England Biolabs). Cas9 mRNA was produced by in vitro transcription of 1 µg template using the mMessage mMachine T3/T7 kit. Capped, polyadenylated Cas9 mRNA was made using the Poly(A) kit (Ambion). The reaction was precipitated using 30 µl lithium chloride solution (Ambion). RNA was eluted in 30 µl nuclease free H_2_O, aliquoted and stored at -80°C until use. A 1 µl sample was run on a 1% agarose gel alongside the RiboRuler High Range ladder (Thermo Scientific) to check for correct size of product (~3kb).

A mixture containing approximately 1 µg Cas9 mRNA and 400 ng/µl of sgRNAs was injected into the animal pole of embryos at the one-cell stage. To assess the effects, high resolution melt analysis (HRMA) was performed using the MeltDoctor^TM^ HRM Mastermix (Life Technologies, 5 µl MeltDoctor^TM^ HRM Mastermix, 1 µl Diluted DNA (20ng/µl; obtained by digesting 24 hour embryos in 20 µg/µl proteinase K at 55˚C for 60 minutes), 0.3 µl forward and reverse primer (10 µM) and 3.4 µl nuclease free H_2_O) under the following conditions: 95°C 10 min, 40 cycles of [95°C 15s, 60°C 1 min], 95°C 10s, 60°C 1 min, 95°C 15s, 60°C 15s on the 7500 Fast Real – Time PCR system (Applied Biosystems). Nucleic acid concentration was measured using a Nanodrop® ND-1000 (Thermoscientific). Concentrations of nucleic acids were measured by the absorbance at 260 nm and the purity of the sample was quantified by the ratio of sample absorbance at 260/280 nm. A lower limit of 1.10 was set as the boundary for an acceptable reading for the 260/280 values due to impurities interfering with the HRMA analysis. During each HRMA reaction, 3 wild type samples were used as controls and each test sample was run as a replicate.

### Active avoidance conditioning

Conditioning was carried out using a two-way chamber, essentially as described in (10), with the addition of a separator, made of matte black cardboard, between the two compartments. The conditioning stimulus (CS), a red LED, was delivered for 8 seconds, while the unconditioned stimulus (US), a 25 V pulse, was delivered for 100 milli-seconds. All fish were genotyped after the assay by sequencing.

### Electrophysiology

Whole cell patch clamp recordings were done as described in (22) from vHb neurons in 6-8 dpf larvae. Briefly, the larvae were anesthetized in 0.01% MS222 and pinned onto a Sylgard (Dow Corning, Midland, MI, United States) dish using fine tungsten wire (California Fine Wire, Grover Beach, CA, United States). The MS222 was then replaced with external saline (composition in mM; 134 NaCl, 2.9 KCl, 1.2 MgCl2, 10 HEPES, 10 Glucose, 2.1 CaCl2, 0.01 D-tubocurare; pH: 7.8; 290 mOsm) and the skin over the head was carefully peeled off to expose the brain. The recording chamber was then transferred to the rig apparatus. All recordings were done in an awake, *in vivo* condition. The cells were observed using a 60× water immersion objective on a compound microscope (Olympus BX61WI). Cells in the ventral most layer of the habenula and on the lateral most side were targeted for recordings as these cells express the Kiss1 receptor. Pipettes of tip diameter 1–1.5 µm and resistance of 12–16 MΩ were pulled with thick walled borosilicate capillaries (1.5 mm OD; 0.86 mm ID; Warner Instruments, Hamden, CT, United States) using a Flaming Brown P-97 pipette puller (Sutter Instruments, Novato, CA, United States). A potassium gluconate based patch internal solution (composition in mM: 115 K gluconate, 15 KCl, 2 MgCl2, 10 HEPES, 10 EGTA, 4 MgATP; pH: 7.2; 290 mOsm) was used for all recordings. Whole cell recordings were acquired using Multiclamp 700B amplifier, Digidata 1440A digitizer and pCLAMP software (Molecular Devices). The data was low pass filtered at 2 kHz using a Bessel filter and sampled at 20 kHz at a gain of 1. Membrane potentials mentioned were not corrected for liquid junction potential which was measured to be +8 mV for the potassium gluconate based internal solution. The following drugs were perfused in the bath: TTX (1 µM, Alomone labs, Israel), K10 (Isca biochemicals, UK) and K234 (Isca biochemicals). Events were detected offline using Clampfit (Molecular Devices). Graphs were plotted using Microsoft Excel. Statistical analysis was done using MATLAB.

## Acknowledgements

We thank Satoshi Ogawa and Ishwar Parhar for the antibodies to kisspeptin1 and the kisspeptin receptor. CL was funded by the A*Star Graduate Academy, under the ARAP scheme. JC was funded by the NUS Graduate School. This work was funded by core funding from IMCB and a Lee Kong Chian School of Medicine, Nanyang Technological University MOE Start-Up Grant to SJ, and core funding from NCBS to VT. We thank Caroline Kibat for performing the antibody label and associated imaging, and Samuel James for help with the behavioural assay.

